# Genetic risk of dementia modifies the impact of obesity on limbic white matter and spatial navigation behavior in cognitively healthy adults

**DOI:** 10.1101/871160

**Authors:** Jilu P. Mole, Fabrizio Fasano, John Evans, Rebecca Sims, Derek A. Hamilton, Emma Kidd, Claudia Metzler-Baddeley

**Author notes:** Corresponding author: Claudia Metzler-Baddeley, CUBRIC, Maindy Road, Cardiff CF24 4HQ. Author contributions: CMB: conceptualization, methodology, formal analysis, writing – original draft preparation, writing – review & editing, visualization, funding acquisition; JPM: investigation, formal analysis, data curation, project administration; RS, EK, DH: Resources; FF, JE: Software.

## Abstract

A family history (FH) of dementia, *APOE*-ε4 genotype, and obesity are major risk factors for developing Alzheimer’s disease but their combined effects on the brain and cognition remain elusive. We tested the hypothesis that these risk factors affect apparent white matter (WM) myelin and cognition including spatial navigation and processing speed in 166 asymptomatic individuals (38-71 years). Microstructure in temporal [fornix, parahippocampal cingulum, uncinate fasciculus], motor and whole-brain WM was assessed with myelin-sensitive indices from quantitative magnetization transfer [macromolecular proton fraction (MPF)] and axon density from diffusion imaging. Individuals with the highest genetic risk (FH+ and *APOE*-ε4) compared to those with FH+ alone showed obesity-related reductions in MPF and axon density in the right parahippocampal cingulum. No effects were present for those without FH. Furthermore, FH modulated obesity-related effects on spatial navigation behaviour. In summary, an individual’s genetic dementia risk influenced the impact of obesity on WM myelin and cognition.

## 1. Introduction

As the world’s population is aging, increasing numbers of people over 65 will develop cognitive impairment due to late onset Alzheimer’s disease (LOAD)^1^. The pathological processes leading to LOAD accumulate over many years^2^ but it remains challenging to reliably identify those individuals at heightened risk before they develop memory impairments. It is therefore important to gain a better understanding of the impact of dementia risk factors on the brain and cognition in cognitive healthy individuals. Identifying reliable biomarkers is pivotal for the development of preventative interventions in the future.

Accumulating evidence suggests that neuroglia damage may be critically involved in the pathogenesis of LOAD^3-9^. Neuroglia are non-neuronal support cells that maintain brain’s homeostasis and are essential for normal synaptic and neuronal activity. In the brain these are myelin-producing oligodendrocytes, microglia immune cells, astrocytes and ependymal cells. According to the myelin hypothesis of LOAD^4^, oligodendrocytes are particularly vulnerable to the impact of aging and insults from genetic and lifestyle risk factors, leading to an accelerated breakdown of axon myelin sheaths with advancing age^3, 10, 11^. Besides providing insulation for axons, myelin regulates saltatory conduction and synaptic plasticity and is therefore critical for the speed of information transfer and the maintenance of healthy brain circuit functions^12-15^. When myelin gets damaged, the brain triggers complex repair mechanisms that may involve Apolipoprotein E (*APOE*) mediated transport of cholesterol^4,16^ as well as beta-secretase enzyme BACE1 mediated neuregulin cleavage to signal oligodendrocytes to myelinate^17-19^. As BACE1 is the key enzyme that initiates the formation of amyloid-β^20^, it has been proposed that amyloid-β plaques may develop as a by-product of myelin repair processes^4^. The myelin model therefore predicts that myelin damage precedes the development of LOAD pathology. Consequently, one may expect accelerated myelin damage in middle-aged and older cognitively healthy individuals at heightened risk of LOAD ^10, 21, 22^. The objective of the present study was to test this hypothesis by investigating the impact of three well-established LOAD risk factors, *APOE*-ε4 genotype, family history of dementia, and central obesity, on myelin-sensitive MRI metrics in brain white matter. Family history was assessed by asking participants whether a first-grade relative was affected by LOAD, vascular dementia, or any other type of dementia.

The protein *APOE* is involved in cholesterol metabolism and lipid homeostasis in the brain^23^. *APOE* has three alleles ε2, ε3 and ε4 and carriage of *APOE*-ε4 is the largest known genetic risk factor of LOAD with ε4 homozygotes having a 14-fold increase in their lifetime risk of developing LOAD compared with *APOE*-ε2 and ε3 carriers ^24, 25^. *APOE* is the main cholesterol transporter in the brain^16^ and the *APOE*-ε4 isoform is thought to have fewer molecules to deliver cholesterol required for myelin and synaptic repair ^16, 26^. It is well-established that *APOE*-ε4 is associated with an earlier onset of LOAD^24, 27^ and a larger burden of amyloid-β plaques^28-32^. This is particularly the case in those individuals with a FH of LOAD^33, 34^. In cognitively healthy individuals, *APOE*-ε4 has been linked to vulnerabilities in spatial navigation^35^, executive function^36, 37^ and processing speed^38^ and both *APOE*-ε4 and FH to impairments in episodic memory^36, 39-42^. Furthermore, both *APOE*-ε4 and FH are associated with changes in brain areas known to be affected in LOAD, such as posterior cingulate, parietal, prefrontal and temporal cortices including hippocampal and parahippocampal regions ^43-48^. For instance, *APOE*-ε4 carriers with a FH of LOAD had larger axial diffusivity in the uncinate fasciculus (UF), a pathway that connects temporal and prefrontal cortex regions^49^. Similarly, baseline differences in white matter microstructure in the fornix, the parahippocampal cingulum (PHC) and the UF were found to be predictive of episodic memory decline over three years in a cohort of asymptomatic older adults enriched with *APOE*-ε4 and FH of LOAD^50^.

Midlife obesity is another established risk factor for developing LOAD (estimated risk ratio of ∼1.4) ^51 52, 53^, particularly in *APOE*-ε4 carriers^54^. Obesity is associated with white matter microstructural differences especially in pathways of the limbic system and those connecting temporal and frontal lobes including the fornix and the cingulum bundle^55^ as well as with impairments in episodic memory^56^, executive functions, and processing speed^57^. A recent study ^58^ reported stronger correlations between obesity state and performance in executive function and memory tests in *APOE*-ε4 carriers than non-carriers suggesting that *APOE* may modulate the effects of obesity on cognition. Thus, preliminary evidence points to complex interplays between *APOE*-ε4, FH, and obesity but there is a lack of studies into their interactive effects on the brain and cognition, and the underlying mechanisms remain unknown.

We therefore investigated the interaction effects between *APOE*, family history of dementia, and Waist-Hip-Ratio, a measure of central obesity, in 166 asymptomatic individuals (38 - 71 years) from the Cardiff Ageing and Risk of Dementia Study (CARDS)^59, 60^ with a focus on white matter microstructure and cognition (Table 1). Based on the myelin model, we hypothesized that risk-related effects would be particularly apparent in myelin-sensitive MRI metrics in temporal lobe pathways known to be susceptible to LOAD, i.e. the fornix, PHC and UF ^61-64^, compared with motor cortico-spinal tracts (CST) and whole brain white matter (WBWM) control regions (Figure 1).

**Table 1.** Summary of demographic, genetic, and lifestyle risk information of participants.^*^Based on waist to hip ratio ≥ 0.9 for males and ≥ 0.85 for females ^112^. Abbreviations: *APOE*, Apolipoprotein-E; FH, Family History of dementia; MMSE = Mini Mental State Exam^110^; NART = National Adult Reading Test^109^; PHQ-9 = Patient Health Questionnaire^111^.

| Mean (SD) | Participants (n = 166) |
| --- | --- |
| Age (in years) | 55.8 (8.2) |
| Females | 56% |
| Years of education | 16.5 (3.3) |
| NART-IQ | 116.7 (6.7) |
| MMSE | 29.1 (1.0) |
| PHQ-9 | 2.6 (2.9) |
| FH+ | 35.1% |
| APOE4+ | 36.3% (4.2% ε4/ε4) |
| Abdominal obesity* | 61.3% |
| Systolic hypertension (≥ 140 mm Hg) | 28% |
| Smokers | 5.4% |
| Diabetes | 1.8% |
| Statins | 7.2% |
| Alcohol units per week | 7.4 (9.3) |
| C-Reactive Protein (lg ng/ml) | 3.1 (0.47) |
| Leptin (lg pg/ml) | 4.1 (0.47) |
| Adiponectin (lg ng/ml) | 4.1 (0.26) |

**Figure 1.**
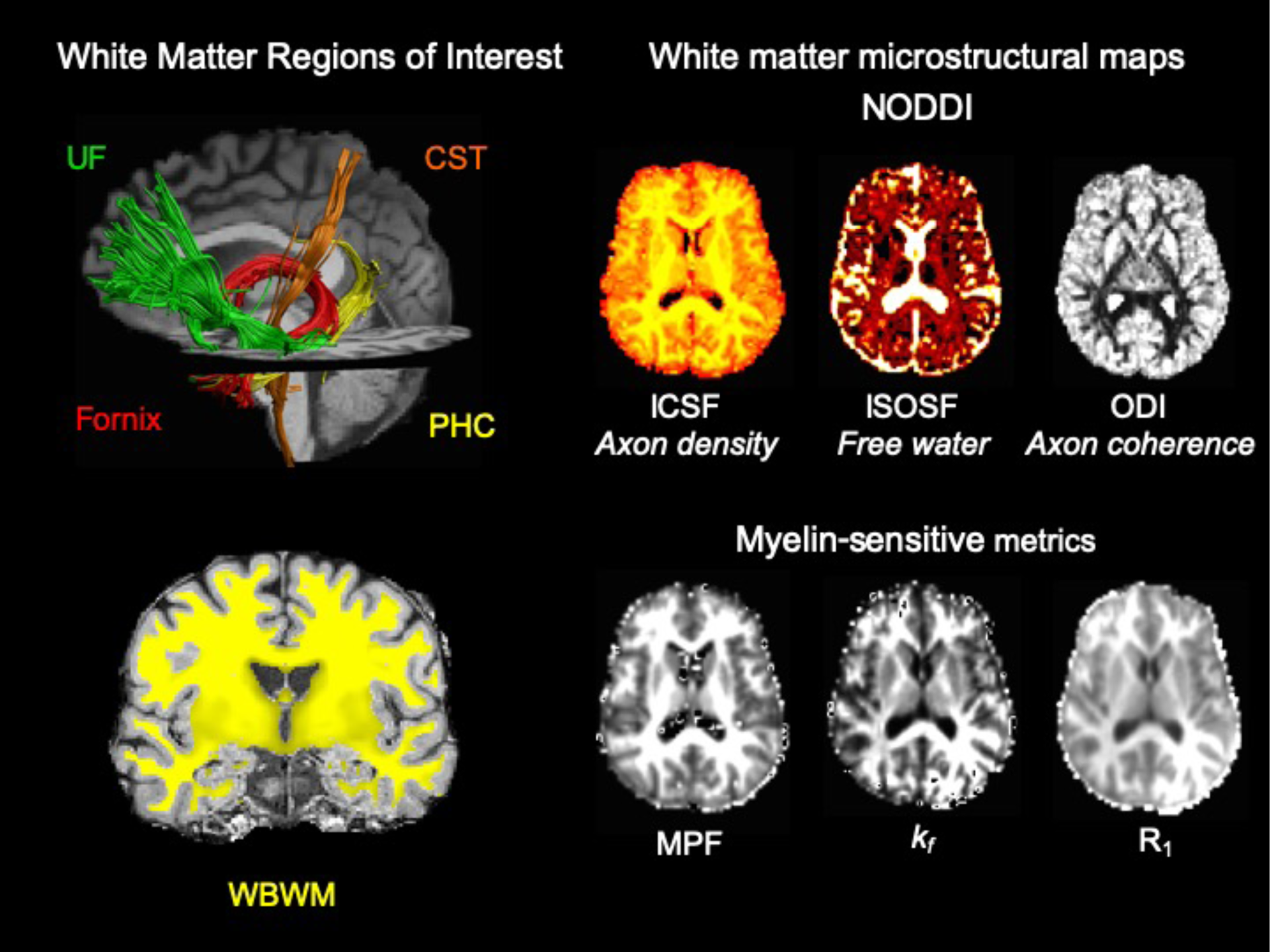
displays the white matter pathways and whole brain white matter region of interest as well as the microstructural maps for one representative participant. Abbreviations: CST, corticospinal tract; ICSF, intracellular signal fraction; ISOSF, isotropic signal fraction; *k*_*f*_, forward exchange rate; MPF, macromolecular proton fraction; NODDI, neurite orientation density and dispersion imaging; ODI, orientation density index; PHC, parahippocampal cingulum; UF, uncinate fasciculus; R_1_, longitudinal relaxation rate; WBWM, whole brain white matter.

Going beyond previously adopted indices from diffusion tensor imaging (DTI)^65, 66,67^, we employed more myelin-sensitive indices from quantitative magnetization transfer (qMT) imaging^68-72^ and T_1_-relaxometry^73, 74^ (longitudinal relaxation rate R_1_). The qMT indices were the macromolecular proton fraction (MPF), known to be highly sensitive to myelin in white matter^72, 75-78^, and the forward exchange rate *k*_*f*_, an index thought to reflect metabolic efficiency that was found to be reduced in LOAD^79^ (Figure 1). In addition, we assessed axon microstructure with dual-shell diffusion-weighted Neurite Orientation Density and Dispersion Imaging (NODDI)^80^. NODDI estimates i) free water contributions to the diffusion signal with the isotropic signal fraction (ISOSF), ii) restricted diffusion contributions with the intracellular signal fraction (ICSF), an estimate of apparent axon density, and iii) axon orientation dispersion (ODI) (Figure 1). Average indices of these microstructural measurements were derived from all white matter pathways and the whole brain region. Based on the myelin model we mainly predicted *APOE*-ε4, family history, and central obesity-related reductions in MPF and R_1_, that may, however, also affect the metabolic index *k*_*f*_ and the axon microstructural indices from NODDI. Furthermore, we investigated the impact of risk factors on a wide range of cognitive domains including episodic memory^65, 81, 82^, spatial navigation^83^, working memory, executive function, and processing speed^84, 85^. As myelin regulates the speed of information transfer in the brain, we expected effects on processing speed such as on response latencies in a virtual Morris Water Maze Task (vMWMT)^86^, that required visual-spatial learning and relied on efficient communication within hippocampal networks. All analyses were controlled for age, sex, and education as these variables are known to affect LOAD risk and allowed the comparison between age and risk-related differences ^87, 88^. Finally, we tested whether differences in blood pressure, plasma markers of inflammation (C-Reactive Protein, Interleukin 8), and adipose tissue dysfunction (leptin/adiponection ratio)^89^ accounted for any effects of risk factors on white matter microstructure and cognition to explore potential contributing mechanisms.

## 2. Results

### 2.1 Multivariate covariance analysis (MANCOVA) of white matter microstructural indices

MANCOVA tested for the effects of *APOE* genotype (ε4+, ε4-), family history (FH+, FH-) and central obesity (WHR+, WHR-) on all microstructural indices (MPF, *k*_*f*_, R_1_, ISOSF, ICSF, ODI) in all white matter pathways (left PHC, right PHC, left UF, right UF, left CST, right CST and fornix) and WBWM regions whilst controlling for age, sex, and years of education.

#### Omnibus effects

Main effects were observed for age [F(48,60) = 2.64, p < 0.001, η_p_^2^= 0.68] and sex [F(48,60) = 2.61, p < 0.001, η_p_^2^ = 0.67]. Interaction effects were present between *APOE* and WHR [F(48,60) = 1.6, p = 0.04, η_p_^2^ = 0.56] and between *APOE*, FH and WHR [F(48,60) = 1.7, p = 0.02, η_p_^2^ = 0.58].

#### Post-hoc effects

Ageing was associated with reductions in MPF, *k*_*f*_, and R_1_ in WBWM, fornix and right UF (Table 2). MPF and *k*_*f*_ reductions were also present in the left UF and bilateral CST respectively. In addition, age-related increases in ISOSF were observed in WBWM and fornix and increases in ODI in the fornix (Table 2, Figure 2A). Women relative to men showed lower *k*_*f*_, ISOSF and ODI in WBWM and the fornix and larger fornix MPF (Table 2, Figure 2B).

**Table 2.** Post-hoc effects on white matter microstructure. ^+^Contrast: centrally obese relative to normal *APOE*-ε4 carriers (*APOE*-ε2/ε4, *APOE*-ε3/ε4, *APOE*-ε4/ε4) with a family history of dementia. ^++^Contrast: centrally obese relative to normal *APOE*-ε4 non-carriers (*APOE*-ε2/ε2, *APOE*-ε2/ε3, *APOE*-ε3/ε3) with a family history. Abbreviations: *APOE*, Apoliopoprotein E; CST, corticospinal tract; FH, family history of dementia; ICSF, intracellular signal fraction; ISOSF, isotropic signal fraction; *k*_*f*_, forward exchange rate; MPF, macromolecular proton fraction; ODI, orientation density index; p_BHadj_, Benjamini-Hochberg adjusted p-value; PHC, parahippocampal cingulum; R_1_, longitudinal relaxation rate; ROI, region of interest; UF, uncinate fasciculus; WM, white matter.

| Effect | ROI | Index | F<sub>(1,107)</sub>-value | p<sub>BHadj</sub> | r | p<sub>BHadj</sub> |
| --- | --- | --- | --- | --- | --- | --- |
| Age | Whole brain WM | MPF | 29.3 | < 0.001 | -0.37 | < 0.001 |
|  |  | ISOSF | 13.7 | < 0.001 | 0.31 | < 0.001 |
|  |  | R<sub>1</sub> | 11.0 | < 0.001 | -0.30 | < 0.001 |
|  |  | k<sub>f</sub> | 10.1 | 0.017 | -0.29 | < 0.001 |
|  | Fornix | MPF | 36.4 | < 0.001 | -0.42 | < 0.001 |
|  |  | ISOSF | 23.9 | < 0.001 | 0.33 | < 0.001 |
|  |  | R<sub>1</sub> | 27.8 | < 0.001 | -0.41 | < 0.001 |
|  |  | k<sub>f</sub> | 29.9 | < 0.001 | -0.41 | < 0.001 |
|  |  | ODI | 7.6 | 0.04 | 0.24 | 0.002 |
|  | L UF | MPF | 9.1 | 0.024 | -0.21 | 0.009 |
|  | R UF | R<sub>1</sub> | 14.6 | < 0.001 | -0.31 | < 0.001 |
|  |  | MPF | 9.0 | 0.023 | -0.21 | 0.008 |
|  |  | k<sub>f</sub> | 9.5 | 0.022 | -0.24 | 0.003 |
|  |  | ICSF | 7.4 | 0.038 | ns |  |
|  | L CST | k<sub>f</sub> | 7.8 | 0.035 | -0.26 | 0.001 |
|  | R CST | k<sub>f</sub> | 14.9 | < 0.001 | -0.36 | < 0.001 |
|  |  |  |  |  | **t-value** |  |
| Sex | Whole brain WM | k<sub>f</sub> | 18.8 | < 0.001 | 3.47 | 0.005 |
|  |  | ISOSF | 11.6 | 0.012 | 2.96 | 0.009 |
|  |  | ODI | 7.3 | 0.043 | 2.9 | 0.009 |
|  | Fornix | ISOSF | 30.7 | < 0.001 | 6.7 | < 0.001 |
|  |  | MPF | 8.9 | 0.024 | 3.1 | 0.006 |
|  | L PHC | k<sub>f</sub> | 9.3 | 0.021 | ns |  |
|  | R PHC | k<sub>f</sub> | 12.7 | 0.011 |  |  |
|  | L UF | k<sub>f</sub> | 9.4 | 0.02 |  |  |
|  | R CST | R<sub>1</sub> | 9.2 | 0.019 |  |  |
| APOE x WHR | R PHC | MPF | 16.4 | < 0.001 | ns |  |
|  |  | ICSF | 7.0 | 0.045 |  |  |
|  | L CST | ODI | 11.7 | 0.01 |  |  |
|  |  | R<sub>1</sub> | 7.0 | 0.04 |  |  |
|  | R CST | MPF | 9.3 | 0.019 |  |  |
|  |  | R<sub>1</sub> | 7.2 | 0.04 |  |  |
| FH x APOE x WHR | Whole brain WM | MPF | 16.9 | < 0.001 | ns |  |
|  | R PHC | MPF | 13.14 | < 0.001 | 3.63<sup>+</sup> | 0.014 |
|  |  |  |  |  | -3.5<sup>++</sup> | 0.014 |
|  |  | ICSF | 11.6 | 0.01 | 3.0<sup>+</sup> | 0.04 |
|  | L CST | MPF | 10.1 | 0.017 | ns |  |
|  | R CST | MPF | 14.9 | < 0.001 |  |  |
|  | L UF | MPF | 12.1 | 0.009 |  |  |
|  | R UF | MPF | 10.7 | 0.009 |  |  |

**Figure 2.**
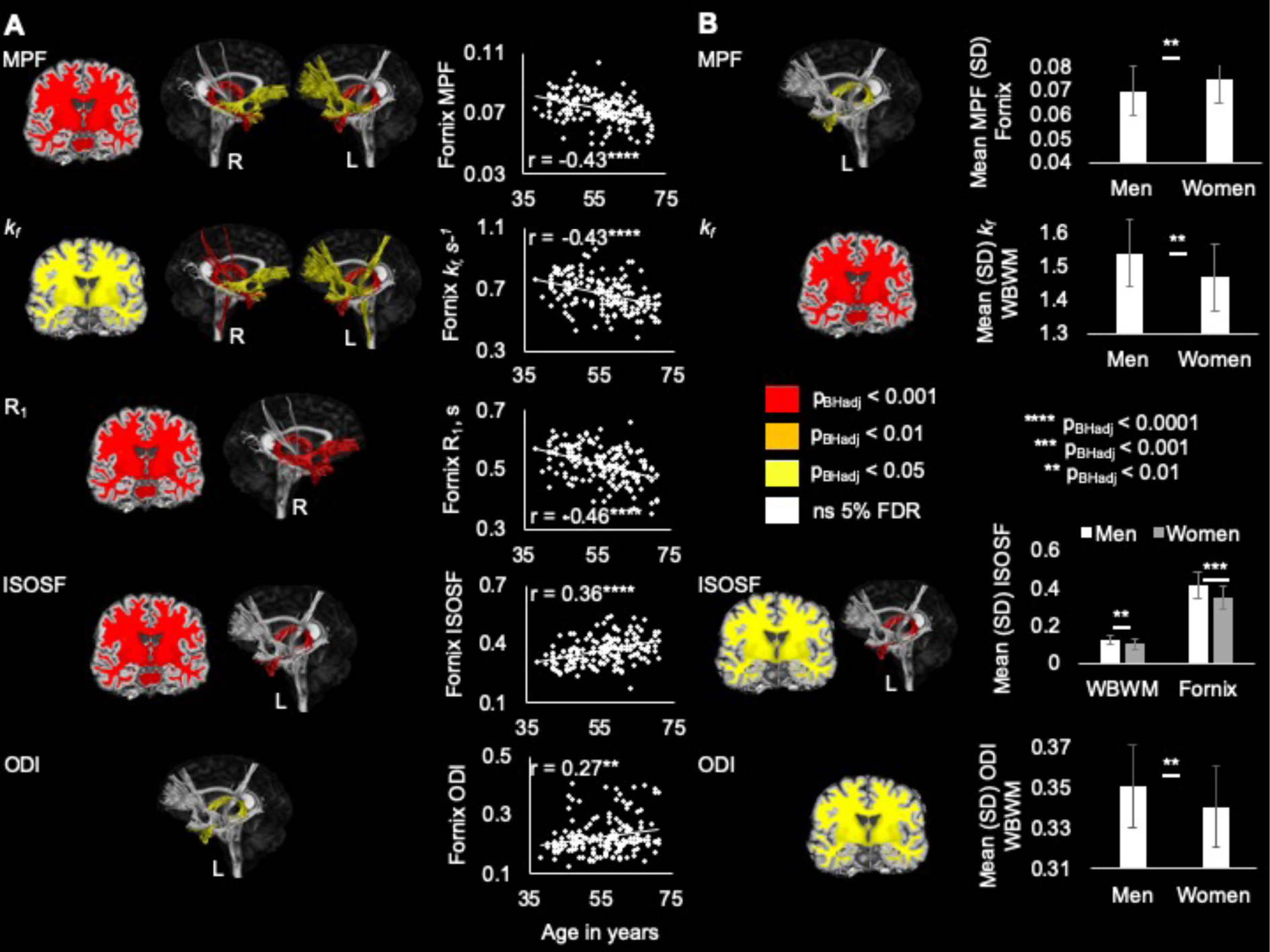
Age (A) and sex (B) effects on white matter microstructure colour-coded according to 5% False Discovery Rate (FDR) corrected p-values. Abbreviations: ISOSF, isotropic signal fraction; *k*_*f*_, forward exchange rate; L, left; MPF, macromolecular proton fraction; ODI, orientation density index; p_BHadj_, p-value corrected for multiple comparison with 5% FDR using the Benjamini-Hochberg adjustement; R, right; SD, standard deviation; WBWM, whole brain white matter.

Two-way interaction effects between *APOE* and WHR were present for right PHC (Figure 3A) and bilateral CST but additional *post-hoc* group comparisons to assess the directions of these effects were non-significant. Three-way interaction effects between FH, *APOE* and WHR were observed for MPF in WBWM, bilateral CST, bilateral UF and right PHC (Figure 3B). For right PHC there was also an effect on ICSF. These three-way interaction effects were followed up by testing for *APOE* and WHR interactions in FH+ and FH-individuals separately (Figure 3C). FH-individuals did not show any interaction effects between *APOE* and WHR [F(7,86) = 0.6, p = 0.72]. In contrast, FH+ individuals exhibited a significant *APOE* x WHR interaction [F(7,43) = 3.7, p = 0.003, η_p_^2^ = 0.38] due to the following pattern: For *APOE*-ε4+ carriers, centrally obese relative to WHR normal individuals had lower MPF and ICSF in the right PHC. The opposite pattern was observed for APOE-ε4-individuals with obese relative to non-obese individuals showing larger MPF in the right PHC (Table 2, Figure 3C).

**Figure 3.**
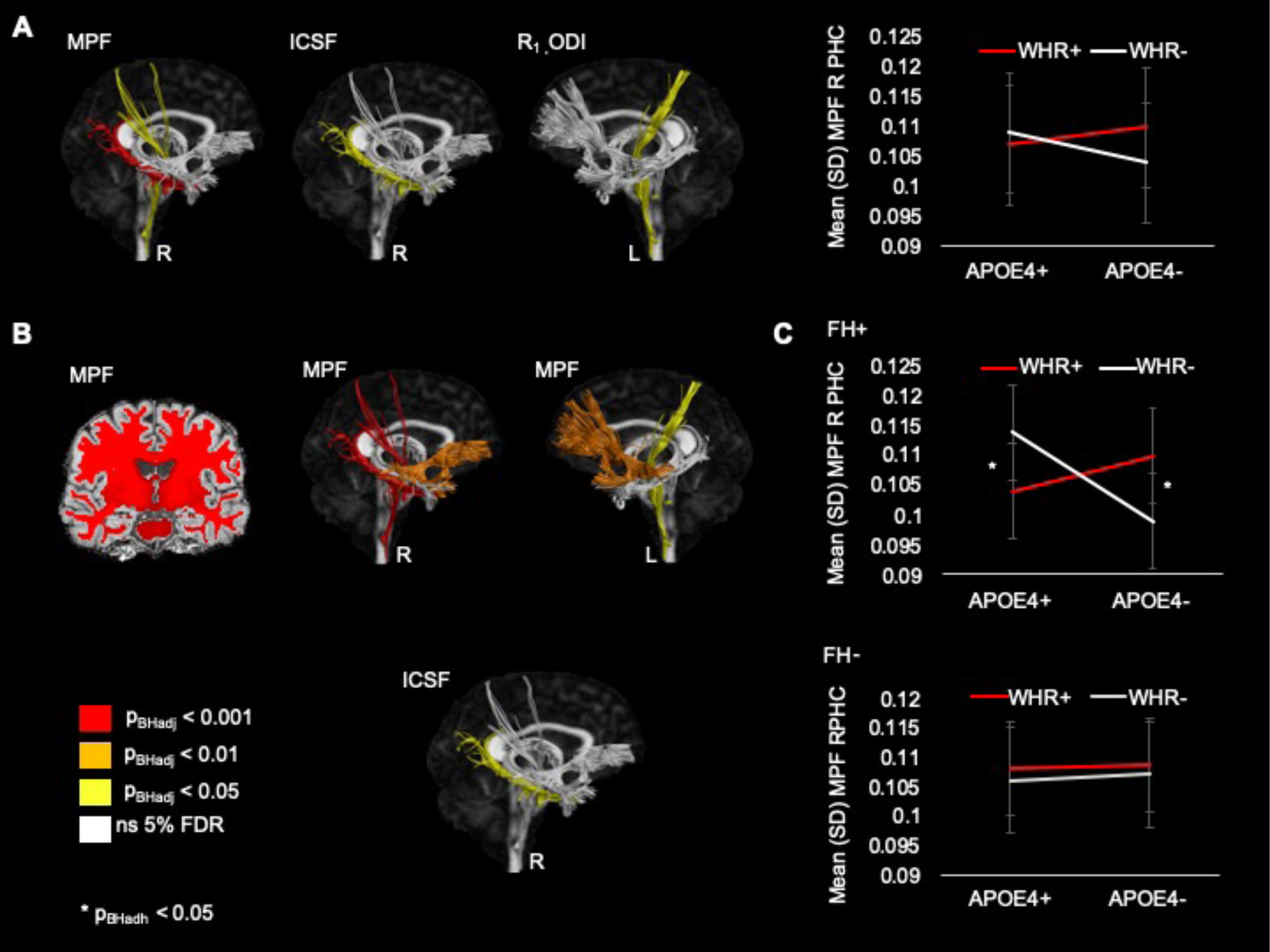
A) Two-way interaction effects between *APOE* genotype and waist hip ratio and B) three-way interaction effects between Family History, *APOE* genotype and Waist Hip Ratio on white matter microstructure colour-coded according to 5% False Discovery Rate (FDR) corrected p-values. Abbreviations: *APOE*, Apolipoprotein E; APOE4+, *APOE*-ε4 carriers (*APOE*-ε2/ε4, *APOE*-ε3/ε4, *APOE*-ε4/ε4); APOE4-, *APOE*-ε4 non-carriers (*APOE*-ε2/ε2, *APOE*-ε2/ε3, *APOE*-ε3/ε3); FH, Family History of dementia; FH+, positive family history; FH-, negative family history; ICSF, intracellular signal fraction; L, left; MPF, macromolecular proton fraction; ODI, orientation density index; p_BHadj_, p-value corrected for multiple comparison with 5% FDR using the Benjamini-Hochberg adjustement; R, right; PHC, parahippocampal cingulum; WHR, Waist Hip Ratio; WHR+, centrally obese; WHR-, WHR in normal range.

We then explored with MANCOVA whether controlling for differences in systolic and diastolic blood pressure (BP), C-Reactive Protein (CRP), Interleukin 8 (IL8), and the leptin/adiponection ratio (LAR) would account for the observed three-way interaction effects for MPF in WBWM, bilateral CST, bilateral UF and right PHC as well as right PHC ICSF. There were significant omnibus effects of systolic [F(7,116) = 4.6, p < 0.001, η_p_^2^ = 0.22] and diastolic blood pressure [F(7,116) = 2.5, p = 0.018, η_p_^2^ = 0.13] but these did not remove the omnibus interaction effect between FH, *APOE* and WHR [F(7,116) = 2.4, p = 0.027, η_p_^2^ = 0.13]. In addition, neither CRP, IL8 or LAR explained the three-way-interaction effects.

*Post-hoc* comparisons revealed significant effects of systolic [F(1,122) = 12.5, p_BHadj_= 0.021, η_p_^2^ = 0.09] and diastolic BP [F(1,122) = 6.8, p_BHadj_ = 0.035, η_p_^2^ = 0.05] on WBWM MPF and of systolic BP on right PHC ICSF [F(1,122) = 6.6, p_BHadj_ = 0.036, η_p_^2^ = 0.05]. Interaction effects between FH, *APOE* and WHR remained significant for right PHC MPF [F(1,122) = 12.0, p_BHadj_ = 0.01, η_p_^2^ = 0.09] and ICSF [F(1,122) = 7.3, p_BHadj_ = 0.03, η_p_^2^ = 0.06] as well as WBWM [F(1,122) = 10.0, p_BHadj_ = 0.01, η_p_^2^ = 0.08] and right UF MPF [F(1,122) = 7.7, p_BHadj_ = 0.03, η_p_^2^ = 0.06].

### 2.2 Exploratory Factor Analysis of cognitive scores

Exploratory Factor Analysis (EFA) was employed to reduce the dimensionality of the cognitive data to seven factors with an Eigenvalue over two, explaining together 49% of the variance in the cognitive data (Table 3). The first factor captured “Verbal Recall” with high loadings on the Rey Auditory Verbal Learning Test (RAVLT). The second factor captured elements of “Motor planning and speed” with high loadings on first move latencies in all conditions and of total latencies in the motor control condition of the virtual Morris Water Maze Task (vMWMT). A third factor captured “Spatial Navigation” behaviour with high loadings on path lengths and total latencies in the hidden platform condition of the vMWMT. The fourth factor captured “Attention Set” with high loadings on the intradimensional subcomponents of the an intra- and extradimensional (IDED) attention set task. The fifth factor captured “Visuospatial Memory” with loadings on immediate and delayed recall of the Rey Complex Figure and the Spatial Span task. The sixth factor captured “Working Memory” capacity due to loadings on Digit Span and Spatial Search. Finally, the seventh “Paired Associated Learning” factor had high loadings on an object object-location paired associate learning task.

**Table 3.**
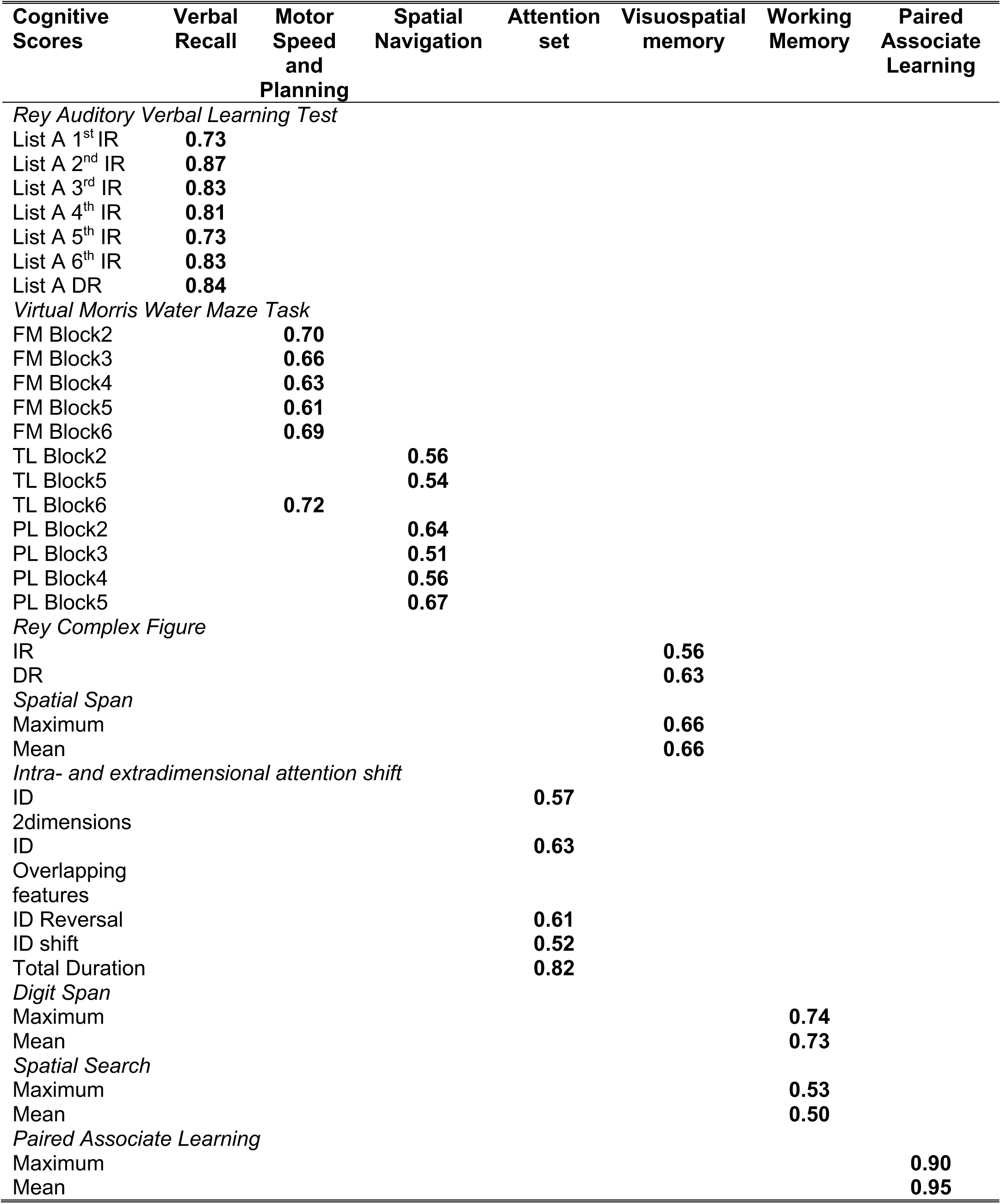
Rotated factor matrix of the exploratory factor analysis within the cognitive data (Rotation methods: Varimax with Kaiser normalization). Loadings > 0.5 are displayed. Abbreviations: DR, delayed recall; ED, extra-dimensional; FM, first move latencies; ID, intra-dimensional; IR, immediate recall; PL, path length; RT, Reaction Times; TL, total latencies.

### 2.3 MANCOVA of cognitive measurements

MANCOVA testing for the effects of *APOE* genotype, FH and WHR on the seven cognitive factors were carried out whilst controlling for age, sex and years of education.

#### Omnibus effects

There was a main effect of age [F(7,105) = 5.6, p < 0.001, η_p_^2^ = 0.27] and an interaction effect between FH and WHR [F(7,105) = 2.7, p = 0.014, η_p_^2^ = 0.15].

#### Post-hoc effects

Age had significant effects on the “Motor Speed and Planning” factor [F(1,111) = 16.9, p_BHadj_ < 0.001, η_p_^2^ = 0.13] and the “Spatial Navigation” factor [F(1,111) = 7.7, p_BHadj_ = 0.03, η_p_^2^ = 0.07] (Table 3) reflecting the fact that older age was associated with larger response latencies and path lengths in the vMWMT (r_Age-MotorSpeed_ = 0.35, p < 0.001; r_Age-SpatialNavigation_ = 0.23, p = 0.019, controlled for sex and education) (Figure 4A). In addition, there was a significant interaction between FH and WHR for the “Spatial Navigation” factor [F(1,111) = 7.6, p_BHadj_ = 0.014, η_p_^2^ = 0.09]. For those with FH, obese individuals performed worse in Spatial Navigation (larger latencies and path lengths) than non-obese individuals whilst those without FH showed the opposite trend (Figure 4B). Controlling for diastolic and systolic BP, CRP, IL8 and LAR did remove this interaction effect [p = 0.2, η_p_^2^ = 0.017] with diastolic BP showing a trend for a contributing effect (p = 0.06).

**Figure 4.**
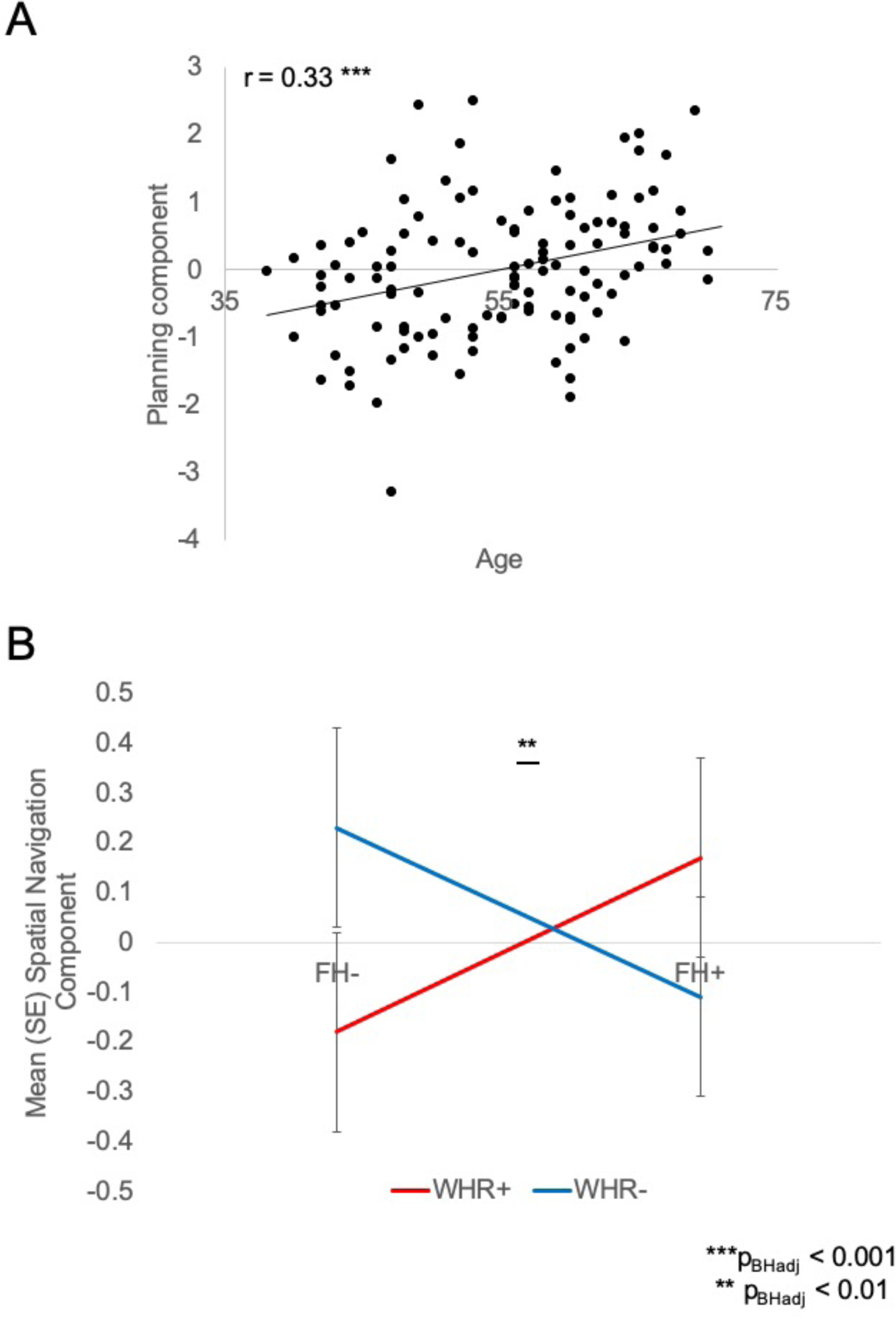
A) Positive correlation between differences in age and the motor planning component. B) Two-way interaction effect between family history of dementia and waist hip ratio and the spatial navigation component. Abbreviations: FH, Family History of dementia; FH+, positive family history; FH-, negative family history; p_BHadj_, p-value corrected for multiple comparison with 5% FDR using the Benjamini-Hochberg adjustement; SE, standard error; WHR, Waist Hip Ratio; WHR+, centrally obese; WHR-, WHR in normal range.

### 2.4 Hierarchical regression analyses testing for white matter-spatial navigation relationships

To find out whether differences in white matter microstructure were predictive of performance differences in the Spatial Navigation Component, hierarchical regression analyses were carried out, that first accounted for the effects of age, sex, years of education before testing for the effects of all microstructural indices in a stepwise fashion. The first model (age, sex, years of education) explained 11% of the variance of the spatial navigation factor [F(3,88) = 3.58, p = 0.02] with significant contributions from age (beta = 0.22, p = 0.03) and years of education (beta = 0.23, p = 0.027). Adding the second model, significantly increased R^2^ to 0.25 [ΔR^2^= 0.15, F(1,87) = 17.1, p < 0.001] with right CST ICSF (beta = −0.39, p < 0.001), years of education (beta = 0.25, p = 0.009) and age (beta = 0.24, p = 0.014) contributing significantly to the final model [F(4,87) = 7.3, p < 0.001].

## 3. Discussion

While family history of dementia, *APOE*-ε4 genotype, and obesity all increase cognitive deficits and LOAD risk individually, only a few studies have investigated their combined effects on the brain and cognition. Based on the myelin model^4^, we predicted interaction effects of these risk factors on myelin-sensitive white matter metrics. Consistent with this hypothesis, we observed three-way interaction effects between family history, *APOE*-ε4, and WHR on the MPF index, known to be highly sensitive to myelin in WM^72, 75-78^. Whilst interaction effects on MPF were observed bilaterally for CST and UF as well as for the WBWM, they were most prominent in the right PHC, a WM pathway, known to be vulnerable in amnestic Mild Cognitive Impairment and LOAD^61-64^. However, no risk effects were found in the fornix or the left PHC, and three-way interactions were only observed for MPF, with the exception of ICSF in the right PHC.

Post-hoc analyses revealed that interactive effects between *APOE* and WHR were dependent on an individual’s family history state. For those individuals with the highest genetic risk, based on *APOE*-ε4 and a positive family history of dementia, obesity was related to lower MPF and ICSF in the right PHC, whilst *APOE*-ε2 and ε3 carriers with a positive family history showed the opposite effect, i.e. larger MPF in obese compared to non-obese individuals. Meanwhile no effects of *APOE* or WHR were observed for individuals without a family history of dementia.

The PHC connects the posterior cingulate and parietal cortices with the medial temporal lobes^90^. Reductions in functional imaging measures, such as glucose metabolism and blood oxygen-level dependent signal, in these regions have been reported in young and middle-aged *APOE*-ε4 carriers and in LOAD^28, 46^ and have been related to altered memory performance^91^. Recently, Chen et al.^92^ applied graph-based network analysis and found reduced global efficiency and functional connectivity in white matter networks as well as reduced functional connectivity in medial temporal lobe regions in older asymptomatic *APOE*-ε4 carriers ^92^. In this study the right parahippocampal gyrus (rPHG) was the only brain region that showed simultaneous functional and structural dysconnectivity, and node efficiency in rPHG mediated *APOE*-ε4-related effects on delayed recall ^92^. Our study expands these findings by providing novel evidence for risk-related microstructural vulnerabilities of the right PHC, that connects those cortical regions with functional abnormalities in *APOE*-ε4 and LOAD.

Furthermore, we also found behavioral evidence for interaction effects between family history and WHR for the spatial navigation factor. This factor captured response latency and path length performance in trials of a virtual Water Maze task that required navigation to a hidden platform. Obese individuals with a positive family history had larger spatial navigation factor scores than normal-weighted individuals, reflecting longer response latencies and path lengths, whilst those without family history showed the opposite trend (Figure 4B). These results complement previous findings of altered navigational behavior in young *APOE*-ε4 carriers^35^ and provide novel evidence for family history-dependent effects of obesity on spatial navigation behavior in middle-aged and older adults. However, similar to other studies in young and middle-aged cohorts ^35, 42^, no overt impairments in episodic memory, motor planning, or attention-executive functioning were associated with risk. Behavioral alterations were specific to spatial navigation performance, known to rely on communications between parietal cortex and the hippocampal formation^93^. Thus, alterations in apparent myelin of the right PHC and spatial navigation performance may reflect the earliest risk-related vulnerabilities in temporal-parietal brain networks prone to LOAD in our cognitively healthy sample.

It should be noted, however, that our regression results suggested that ICSF differences in the right CST rather than in the right PHC contributed significantly to spatial navigation performance. This may reflect the fact that the vMWMT had a large motor component as participants had to navigate through the virtual pool by pressing buttons on the keyboard. Furthermore, right CST also showed interaction effects between family history, *APOE* and WHR on MPF. CST contributions to spatial navigation are therefore consistent with the view that risk-related alterations in WM microstructure affect cognitive performance.

It is also noteworthy that, in contrast to age and sex, we did not observe any main effects of risk on WM microstructure or cognition. This pattern of results suggests complex interplays between an individual’s genetic make-up and obesity. Besides *APOE* genotype, genome wide association studies (GWAS) have identified a number of genetic risk loci for LOAD, involved in lipid metabolism, immunity as well as tau binding and amyloid precursor protein processing ^94^. As self-report of parental history of dementia is likely to represent the presence of some of these genetic risk variants^95^, our effects most likely reflect interactions between *APOE*, obesity and an individuals’ polygenic risk for dementia.

The precise mechanisms underpinning these interaction effects require future studies into the role of a number of metabolic pathways related to lipids, immunity, and vascular function, as well as glucose and insulin resistance. Here, we explored the potential moderating effects of systolic and diastolic BP, of plasma markers of inflammation with CRP and IL8, and of adipose tissue dysfunction with the Leptin/Adiponectin Ratio, that may reflect insulin resistance and/or low-grade inflammation^89^. Differences in BP had an impact on WBWM MPF and right PHC ICSF but controlling for these variables did not fully account for the observed interaction in microstructure. In contrast, controlling for the variables removed the interaction effect between FH and WHR on the spatial navigation factor. This was likely driven by diastolic BP as the only variable that showed a trend for a contributing effect (p = 0.06). Hypertension is a known risk factor of cognitive impairment^96^ and hence may contribute to the observed interactions.

As myelin synthesis and maintenance are highly sensitive to cholesterol concentrations, and both *APOE*-ε4 and obesity are associated with hypercholesterolemia^51^, it is also plausible that differences in cholesterol metabolism pathways may have contributed to our findings. Although cholesterol concentrations in the brain are largely independent of peripheral cholesterol levels, there is evidence of positive associations between WM microstructure and serum cholesterol^97^ and cognition^98, 99^. In contrast, hypercholesterolemia in older adults or the presence of other cardiovascular risk factors is associated with lower WM microstructure and increased risk of LOAD^99-101^. Positive associations between WM microstructure and allele dosage of the cholesterol gene CETP (cholesteryl ester transfer protein), which can influence the *APOE*-related risk of LOAD^102^, were also observed for young individuals whilst the opposite pattern was found for older adults^97^. Similarly, Birdsill et al.^103^ reported greater waist circumference to be associated with larger WM fractional anisotropy and lower diffusivities in middle-aged community-dwelling individuals whilst other studies found obesity-related reductions of WM microstructure^55, 104^. In summary, these results together with the findings presented here, suggest that the direction of obesity effects on WM microstructure may be dependent on genetic (polygenic risk, *APOE*), cardiovascular and demographic (age, sex) factors that need to be taken into account in future obesity imaging studies.

In light of this evidence one may speculate that *APOE*-ε4 in combination with other polygenic risk factors potentially captured, at least in part, by family history, interacts adversely with obesity and triggers accelerated myelin damage and impaired myelin repair. Myelin decline may particularly affect obese *APOE*-ε4 carriers with a family history, due to a combination of metabolic, vascular, and inflammatory processes that cause myelin damage in the presence of inefficient cholesterol transport for myelin repair. In contrast, obese individuals without genetic risk of LOAD and other cardiovascular or demographic risk factors, may exhibit larger WM myelination due to hypercholesterolemia that may be accompanied by superior cognitive performance. Such complex interactions between genetic factors and lipid pathways may potentially hold an explanation for the “obesity paradox”, that refers to studies in the literature that have reported protective effects of obesity^105^.

Our study also provides insights into differential effects of aging, sex and risk factors on white matter microstructure and cognition. Aging was associated with widespread reductions in MPF, *k*_*f*_, and R_1_ and with increases in ISOSF and ODI particularly in the fornix and WBWM but also in UF and CST. The only pathways not affected by age were left and right PHC. Similarly, women relative to men showed lower *k*_*f*_, ISOSF and ODI in WBWM and the fornix as well as larger MPF in the fornix but no differences in PHC. Previously, we reported negative correlations between central obesity measures, including estimates of abdominal visceral fat, and qMT indices (MPF, *k*_*f*_), that were particularly prominent in the fornix tract without any effects of family history or *APOE*^60^. Thus, while interaction effects between family history, *APOE* and WHR were observed for MPF (and ICSF) and were most prominent in the PHC, they were absent in the fornix. In contrast, age and sex affected all microstructural indices most prominently in the fornix but not in the PHC. Together this pattern of results suggests that genetic-related risk factors (*APOE* and family history) affect medial temporal lobe input (PHC) but not output (fornix) structures, whilst age, sex and obesity affect output (fornix) but not input (PHC) structures. As *APOE* and family history are associated with earlier and larger amyloid and tau burden^33, 106^, and LOAD pathology is known to spread from the entorhinal cortices, that are adjacent to the parahippocampal gyrus, into the medial temporal lobes^107, 108^, this pattern may reflect tissue vulnerabilities that may precede the development of LOAD pathology.

Thus, the pivotal question concerns whether the observed cross-sectional risk effects on MPF and spatial navigation are predictive of accelerated development of LOAD pathology and cognitive and neuronal decline. A future prospective longitudinal follow-up study of the CARDS participants is required to answer this question. This study should also include GWAS based information about individuals’ polygenic risk of LOAD, and a more comprehensive investigation of inflammatory, vascular, lipid and glucose/insulin moderators to inform about potential mechanisms.

## 4. Materials and Methods

Ethical approval for this study was granted by the School of Psychology Research Ethics Committee at Cardiff University (EC.14.09.09.3843R2). All participants provided written informed consent in accordance with the Declaration of Helsinki.

### 4.1 Participants

Participants between the age of 35 and 75 years were recruited for the Cardiff Ageing and Risk of Dementia Study (CARDS) ^59, 60^ *via* Cardiff University community panels and notice boards as well as *via* internet and local poster advertisements. Participants had a good command of the English language and were without a history of neurological and/or psychiatric disease, head injury with loss of consciousness, drug or alcohol dependency, high risk cardio-embolic source or known significant large-vessel disease. N = 166 CARDS volunteers that fulfilled MRI screening criteria underwent scanning at the Cardiff University Brain Research Imaging Centre (CUBRIC). Table 1 summarises information about demographics, cognition, and genetic and lifestyle dementia risk for these 166 participants (as described in ^59, 60^). Participants’ intellectual function was assessed with the National Adult Reading Test (NART) ^109^, cognitive impairment was screened for with the Mini Mental State Exam (MMSE) ^110^ and depression with the PHQ-9 ^111^. All participants had an MMSE ≥ 26. Eight participants scored ≥ 10 in the PHQ-9 suggesting moderate levels of depression but no participant was severely depressed.

### 4.2 Assessment of genetic risk factors

Saliva samples were collected with the Genotek Oragene-DNA kit (OG-500) for DNA extraction and *APOE* genotyping. *APOE* genotypes ε2, ε3 and ε4 were determined by TaqMan genotyping of single nucleotide polymorphism (SNP) rs7412 and KASP genotyping of SNP rs429358. Genotyping was successful in 165 of the 166 participants who underwent an MRI scan. Participants also self-reported their family history of dementia, i.e. whether a first-grade relative was affected by Alzheimer’s disease, vascular dementia or any other type of dementia. 30.5% (n = 18) of individuals with a positive family history were *APOE-*ε4 carriers compared with 43% (n = 46) amongst individuals with a negative family history.

### 4.2 Assessment of metabolic syndrome-related risk factors

Central obesity was assessed with WHR following the World Health Organisation’s recommended protocol for measuring waist and hip circumference ^112^. Obesity was defined as a WHR ≥ 0.9 for males and ≥ 0.85 for females. Resting systolic and diastolic blood pressure (BP) were measured with a digital blood pressure monitor (Model UA-631; A&D Medical, Tokyo, Japan) whilst participants were comfortably seated with their arm supported on a pillow. The mean of three BP readings was calculated. Other metabolic risk factors, including diabetes mellitus, high levels of blood cholesterol controlled with statin medication, history of smoking, and weekly alcohol intake were self-reported by participants in a medical history questionnaire ^113^ (Table 1). As there were relatively few diabetics, smokers, and individuals on statins these variables were not included in the statistical analyses.

### 4.3 Blood plasma analysis

Venous fasting blood samples were drawn into 9ml heparin coated plasma tubes after 12 hours overnight fasting and were centrifuged for 10 minutes at 2,000xg within 60 minutes from blood collection. Plasma samples were transferred into 0.5 ml polypropylene microtubes and stored in a freezer at −80°C. Circulating levels of high-sensitivity C-Reactive Protein (CRP) in mg/dL were assayed using a human CRP Quantikine enzyme-linked immunosorbent assay (ELISA) kit (R & D Systems, Minneapolis, USA). Leptin concentrations in pg/ml were determined with the DRP300 Quantikine ELISA kit (R & D Systems) and adiponectin in ng/ml with the human total adiponectin/Acrp30 Quantitkine ELISA kit (R & D Systems) and leptin/adiponectin ratios for each participant were calculated. Interleukin IL-8 levels in pg/mL were determined using a high sensitivity CXCL8/INTERLEUKIN-8 Quantikine ELISA kit (R & D Systems).

### 4.2 Cognitive assessment

For a summary of all cognitive scores see Table 3. Immediate and delayed (30 minutes) verbal and visual recall were assessed with the Rey Auditory Verbal Learning Test (RAVLT) ^81, 82^ and the complex Rey figure^65^. Short term topographical memory was measured with the Four Mountains Test ^114^. Spatial navigation was assessed with a virtual Morris Water Maze Task (vMWMT)^83^ that required participants to find a hidden platform in a water pool. This task was comprised of six blocks of four trials each. The first block was a practise block to familiarise participants with the task. Blocks 2-5 were the experimental blocks with a hidden platform the participant had to navigate to. Block 6 was a motor control condition, where participants navigated to a visible platform. Outcome measures for the vMWMT were mean total latencies, first move latencies, and total path lengths for each block. Working memory capacity and executive functions were assessed with computerised tests from the Cambridge Brain Sciences battery ^84, 85^. Working memory capacity was tested with digit and spatial span, distractor suppression with an adapted version of the Stroop test (Double-Trouble), problem solving with a version of the Tower of London task (the Tree task), abstract reasoning with grammatical reasoning and the odd-one-out task, and the ability to manipulate and organize spatial information with a self-ordered spatial span task. In addition, participants performed an intra- and extradimenstional shift task, a choice-reaction time task, and an object-location paired-associate learning (PAL). Outcome measures for the Cambridge Brain Sciences tasks were responses latencies and mean and maximum number of correct responses.

#### MRI data acquisition

The following MRI data were acquired on a 3T MAGNETOM Prisma clinical scanner (Siemens Healthcare, Erlangen, Germany). T_1_-weighted images were acquired with a three-dimension (3D) magnetization-prepared rapid gradient-echo (MP-RAGE) sequence (256 × 256 acquisition matrix, TR = 2300 ms, TE = 3.06 ms, TI = 850ms, flip angle θ = 9°, 176 slices, 1mm slice thickness, FOV = 256 mm and acquisition time of ∼ 6 min).

Microstructural data were acquired with the following sequences. High Angular Resolution Diffusion Imaging (HARDI)^115^ data (2 × 2 × 2 mm voxel) were collected with a spin-echo echo-planar dual shell HARDI sequence with diffusion encoded along 90 isotropically distributed orientations ^116^ (30 directions at b-value = 1200 s/mm^2^ and 60 directions at b-value = 2400 s/mm^2^) and six non-diffusion weighted scans with dynamic field correction and the following parameters: TR = 9400ms, TE = 67ms, 80 slices, 2 mm slice thickness, FOV = 256 × 256 × 160 mm, GRAPPA acceleration factor = 2 and acquisition time of ∼15 min.

Quantitative magnetization transfer weighted imaging (qMT) data were acquired with an optimized 3D MT-weighted gradient-recalled-echo sequence ^68^ to obtain magnetization transfer-weighted data with the following parameters: TR = 32 ms, TE = 2.46 ms; Gaussian MT pulses, duration t = 12.8 ms; FA = 5°; FOV = 24 cm, 2.5 × 2.5 × 2.5 mm^3^ resolution. The following off-resonance irradiation frequencies (Θ) and their corresponding saturation pulse nominal flip angles (ΔSAT) for the 11 MT-weighted images were optimized using Cramer-Rao lower bound optimization: Θ = [1000 Hz, 1000 Hz, 2750 Hz, 2768 Hz, 2790 Hz, 2890 Hz, 1000 Hz, 1000 Hz, 12060 Hz, 47180 Hz, 56360 Hz] and their corresponding ΔSAT values = [332°, 333°, 628°, 628°, 628°, 628°, 628°, 628°, 628°, 628°, 332°]. The longitudinal relaxation time, T_1_, of the system was estimated by acquiring three 3D gradient recalled echo sequence (GRE) volumes with three different flip angles (θ = 3°,7°,15°) using the same acquisition parameters as used in the MT-weighted sequence (TR = 32 ms, TE = 2.46 ms, FOV = 24 cm, 2.5 × 2.5 × 2.5 mm^3^ resolution). Data for computing the static magnetic field (B_0_) were collected using two 3D GRE volumes with different echo-times (TE = 4.92 ms and 7.38 ms respectively; TR= 330ms; FOV= 240 mm; slice thickness 2.5 mm) ^117^.

### 4.5 HARDI and qMT data processing

A detailed description of the microstructural data processing was provided in Metzler-Baddeley et al. ^59, 60^. The dual-shell HARDI data were split and b = 1200 and 2400 s/mm^2^ data were corrected separately for distortions induced by the diffusion-weighted gradients and motion artifacts with appropriate reorientation of the encoding vectors ^118^ in ExploreDTI (Version 4.8.3) ^119^. EPI-induced geometrical distortions were corrected by warping the diffusion-weighted image volumes to the T_1_ –weighted anatomical images ^120^. After preprocessing, the NODDI model ^80^ was fitted to the HARDI data with the fast, linear model fitting algorithms of the Accelerated Microstructure Imaging via Convex Optimization (AMICO) framework ^121^ to gain ISOSF, ICSF, and ODI maps.

Using Elastix ^122^, MT-weighted GRE volumes were co-registered to the MT-volume with the most contrast using a rigid body (6 degrees of freedom) registration to correct for inter-scan motion. Data from the 11 MT-weighted GRE images and T_1_-maps were fitted by a two-pool model using the Ramani pulsed-MT approximation ^123^. This approximation provided MPF and *k*_*f*_ maps. MPF maps were thresholded to an upper intensity limit of 0.3 and *k*_*f*_ maps to an upper limit of 3.0 using the FMRIB’s fslmaths imaging calculator to remove voxels with noise-only data.

All image modality maps and region of interest masks were spatially aligned to the T_1_-weighted anatomical volume as reference image with linear affine registration (12 degrees of freedom) using FMRIB’s Linear Image Registration Tool (FLIRT).

### 4.6 Tractography

The RESDORE algorithm ^124^ was applied to identify outliers, followed by whole brain tractography with the damped Richardson-Lucy algorithm (dRL) ^125^ on the 60 direction, b = 2400 s/mm^2^ HARDI data for each dataset in single-subject native space using in house software ^124^ coded in MATLAB (the MathWorks, Natick, MA). To reconstruct fibre tracts, dRL fibre orientation density functions (fODFs) were estimated at the centre of each image voxel. Seed points were positioned at the vertices of a 2×2×2 mm grid superimposed over the image. The tracking algorithm interpolated local fODF estimates at each seed point and then propagated 0.5mm along orientations of each fODF lobe above a threshold on peak amplitude of 0.05. Individual streamlines were subsequently propagated by interpolating the fODF at their new location and propagating 0.5mm along the minimally subtending fODF peak. This process was repeated until the minimally subtending peak magnitude fell below 0.05 or the change of direction between successive 0.5mm steps exceeded an angle of 45°. Tracking was then repeated in the opposite direction from the initial seed point. Streamlines whose lengths were outside a range of 10mm to 500mm were discarded.

The fornix, parahippocampal cinguli (PHC), uncinate fasciculus (UF) and corticospinal (CST) pathways were reconstructed with an in-house automated segmentation method based on principal component analysis (PCA) of streamline shape ^126^. This procedure involves the manual reconstruction of a set of tracts that are then used to train a PCA model of candidate streamline shape and location. Twenty datasets were randomly selected as training data. Tracts were reconstructed by manually applying waypoint region of interest (ROI) gates (“AND”, “OR” and “NOT” gates following Boolean logic) to isolate specific tracts from the whole brain tractography data. ROIs were placed in HARDI data native space on colour-coded fiber orientation maps (Pajevic & Pierpaoli, 1999) in ExploreDTI following previously published protocols ^62, 113, 127, 128^. The trained PCA shape models were then applied to all datasets: candidate streamlines were selected from the whole volume tractography as those bridging the gap between estimated end points of the candidate tracts. Spurious streamlines were excluded by means of a shape comparison with the trained PCA model. All automatic tract reconstructions underwent quality control through visual inspection and any remaining spurious fibers that were not consistent with the tract anatomy were removed from the reconstruction where necessary. Mean values of all qMT and NODDI indices were extracted for all white matter pathways.

### 4.7 Whole brain white matter segmentation

Whole brain white matter masks were automatically segmented from T_1_-weighted images with the Freesurfer image analysis suite (version 5.3), which is documented online (https://surfer.nmr.mgh.harvard.edu/). Whole brain white matter masks were thresholded to exclude ventricular cerebrospinal fluid spaces from the mask. Mean values of all qMT and NODDI indices were extracted for the whole brain white matter mask.

### 4.8 Statistical analysis

Statistical analyses were conducted in SPSS version 20^129^. All microstructural data were examined for outliers defined as above or below three times of the interquartile range (75th percentile value - 25th percentile value). This led to an exclusion of 2.3% of the microstructural and 2.4% of the cognitive data.

Separate multivariate analyses of covariance (MANCOVA) were carried out to test for the effects of *APOE* genotype (ε4+, ε4-), FH (FH+, FH-) and central obesity (WHR+, WHR-) on i) all microstructural indices (MPF, *k*_*f*_, R_1_, ISOSF, ICSF, ODI) in all white matter pathways (left PHC, right PHC, left UF, right UF, left CST, right CST and fornix) and WBWM regions and ii) on all cognitive factors (see below). These analyses included age, sex, and years of education as covariates. Significant omnibus effects were further investigated with post-hoc comparisons across all outcome measures. These were corrected for multiple comparisons with a False Discovery Rate (FDR) of 5% using the Benjamini-Hochberg procedure ^130^ (p_BHadj_). Information about effects sizes is provided with the partial eta squared index η_p_^2^.

The dimensionality of the cognitive data was reduced with exploratory factor analyses (EFA) with unweighted least squares and orthogonal varimax rotation with a maximum of 5000 iterations for conversion. Variables with communalities < 0.4 were excluded from the final EFA^131^. After inspection of Cattell’s scree plot^132^, factors with an eigenvalue exceeding 2 were extracted and factor loadings exceeding 0.5 were considered for the interpretation of the factors. The resulting seven cognitive factors (Table 3) were then entered as dependent variables into the above described MANCOVA models as well as into linear hierarchical regression models, which firstly tested for the effects of age, sex, and years of education and secondly for the effects of all WM microstructural indices in a stepwise fashion.

## Acknowledgements

This research was funded by a Research Fellowship awarded to CM-B from the Alzheimer’s Society and the BRACE Alzheimer’s Charity (grant ref: 208). We would like to thank Erika Leonaviciute, Peter Hobden and Sonya Foley-Bozorgzad for their assistance with MRI data acquisition and processing and Rosie Dwyer, Samantha Collins, Abbie Stark, and Emma Blenkinsop for their assistance with the collection and scoring of the cognitive and health data. We would also like to thank Rhodri Thomas for his assistance with the APOE genotyping of the saliva samples and Benjamin Ertefai for his assistance with the blood plasma analyses.

